# miRNA profiling of primate cervicovaginal lavage and extracellular vesicles reveals miR-186-5p as a potential retroviral restriction factor in macrophages

**DOI:** 10.1101/263947

**Authors:** Zezhou Zhao, Dillon C. Muth, Kathleen Mulka, Zhaohao Liao, Bonita H. Powell, Grace V. Hancock, Kelly A. Metcalf Pate, Kenneth W. Witwer

## Abstract

The goal of this study was to characterize extracellular vesicles (EVs) and miRNAs of primate cervicovaginal lavage (CVL) during the menstrual cycle and simian immunodeficiency virus (SIV) infection, and to determine if differentially regulated CVL miRNAs might influence retrovirus replication. CVL and peripheral blood were collected from SIV-infected and uninfected macaques. EVs were enriched by stepped ultracentrifugation and characterized thoroughly. miRNA profiles were assessed with a medium-throughput stem-loop/hydrolysis probe qPCR platform and validated by single qPCR assays. Hormone cycling was abnormal in infected subjects, but EV concentration correlated with progesterone concentration in uninfected subjects. miRNAs were present predominantly in the EV-depleted CVL supernatant. Only a small number of CVL miRNAs were found to vary during the menstrual cycle or SIV infection. Among them was miR-186-5p, which was depleted in retroviral infection. In experiments with infected macrophages in vitro, this miRNA inhibited HIV replication. These results provide further evidence for the potential of EVs and small RNAs as biomarkers or effectors of disease processes in the reproductive tract.

## 1. Introduction

The cervicovaginal canal is a potential source of biological markers for forensics investigations ^1–4^, reproductive tract cancers ^5–7^, and infections ^8–10^. Cervicovaginal secretions may be collected by swab, tampon, or other methods, or secretion components may be liberated by a buffered wash solution and collected as cervicovaginal lavage (CVL). In addition to utility as biomarkers, constituents of cervicovaginal secretions, including proteins, certain microbes, and metabolites, exert function, for example by playing protective roles in wound healing ^11^ and against HIV-1 infection ^12–22^. A large and important body of work has thus examined biomarker potential and functional roles of numerous entities in the cervicovaginal compartment.

Compared with secreted proteins, metabolites, and the microbiome, however, several components of cervicovaginal fluids are less well understood, including extracellular RNAs (exRNAs) and their carriers, such as extracellular vesicles (EVs) and ribonucleoprotein complexes (exRNPs). EVs are potential regulators of cell behavior in paracrine and endocrine fashion due to their reported abilities to transfer proteins, nucleic acids, sugars, and lipids between cells ^23^. EVs comprise a wide array of double-leaflet membrane extracellular particles, including those of endosomal and cell-surface origin^24,25^, and range in diameter from 30 nm to well over one micron (large oncosomes) ^26^. EV macromolecular composition tends to reflect, but is not necessarily identical to, that of the cell of origin ^27^. EVs have been isolated from most cells, as well as biological fluids ^23,28^, including cervicovaginal secretions of humans ^29^ and rhesus macaques ^30^.

microRNAs (miRNAs) are one of the most studied classes of exRNA. These noncoding RNAs average 22 nucleotides in length and, in some cases, fine-tune the expression of target transcripts ^31,32^. Released from cells by several routes, miRNAs are among the most frequently examined biomarker candidates in biofluids and, along with some other RNAs, are reported to be transmitted via EVs ^33–36^. miRNAs are found not only in EVs, but also in free Argonaute-containing protein complexes; the latter may outnumber the former, at least in blood ^37,38^. Many miRNAs are also highly conserved ^32^, and abundant species typically have 100% identity in humans and nonhuman primates ^39^. (For this reason, we will refer to hsa-(*Homo sapiens*) and mml-(*Macaca mulatta*) miRNAs without the species designation unless otherwise warranted by sequence disparity.) While miRNAs have been profiled in cervicovaginal secretions and menstrual blood, mostly in the forensics setting ^4,40,41^, their associations with EV and exRNP fractions require further study. A recent publication reported that EVs from healthy vaginal secretions inhibited HIV-1 infection ^29^. Another report found that CVL EVs (styled “exosomes”) were present at higher concentrations in cervical cancer, and that two miRNAs were also upregulated ^5^. Our laboratory described a reduction of CVL EVs in a severe endometriosis case compared with reproductively healthy primates ^30^. However, our study, along with others, was limited by the absence of molecular profiling of EV cargo ^30^.

Here, we performed targeted miRNA profiling of EV-enriched and -depleted fractions of CVL and vaginal secretions collected from healthy and retrovirus-infected rhesus macaques. We queried how CVL EVs and miRNAs are affected by the menstrual cycle, an important potential confounder of biomarker studies ^42^. Similarly, we assessed possible associations with simian immunodeficiency virus (SIV) infection. We report an association of miR-186 levels with SIV infection and find that this miRNA also appears to have antiretroviral effects in HIV-infected macrophages. These studies provide baseline information for easily accessed CVL markers including EVs and miRNAs that may become useful tools in the clinic.

## 2. Materials and Methods

### Sample Collection

CVL and whole blood samples were collected weekly for five weeks from two uninfected (control) and four SIVmac251-infected (infected) rhesus macaques (*Macaca mulatta*) as previously described ^30^. All macaques were negative for simian T-cell leukemia virus and simian type D retrovirus and were inoculated intravenously. Animals were sedated with ketamine at a dose of 7-10 mg/kg prior to all procedures. CVL was performed by washing the cervicovaginal cavity with 3 mL of phosphate buffered saline (PBS, Thermo Fisher Scientific, Waltham, MA, USA. Cat #: 14190-144) directed into the cervicovaginal canal and re-aspirated using the same syringe. Materials and procedures for sample collection are depicted in Supplemental Figure 1. Volumes of CVL yield across collection dates were documented in Supplemental Table 1. Whole blood (3 mL) was collected by venipuncture into syringes containing acid citrate dextrose solution (ACD) (Sigma Aldrich, St. Louis, MO, USA. Cat #: C3821).

**Figure 1.**
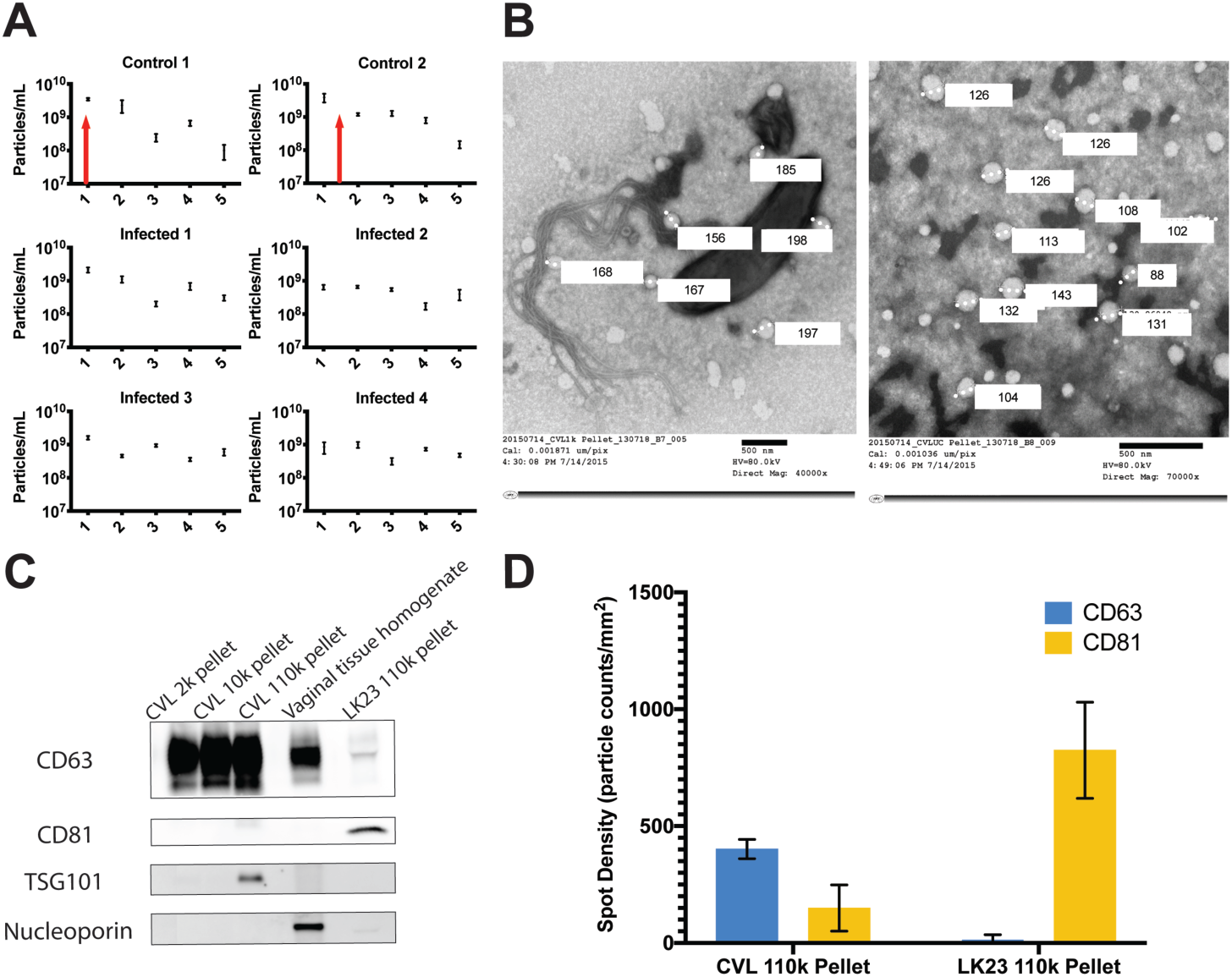
EV composition during the menstrual cycle. **A)** Nanoparticle concentrations of CVL ultracentrifuge (UC) pellets monitored weekly over five weeks for two SIV-negative (“control”) and four SIV-infected rhesus macaques. Red arrows indicate time of ovulation for 2 control animals, which were absent for SIV infected animals. **B**) Transmission electron micrographs of CVL pellets from the 10,000× g pellet **(left)** and 110,000× g pellet **(right)** confirm presence of bacteria and EVs/EV-like particles, with several respective diameters indicated. **C)** Western blot analysis suggests enrichment of EV markers CD63, CD81, and TSG101 in 110k pellet fraction of CVL from uninfected animals. Vaginal tissue homogenate and dendritic cell (DC, LK23) 110k pellet controls were also positive for CD63 and CD81. Nuclear marker nucleoporin was detected in tissue homogenate but not in putative EV samples. **D)** SP-IRIS confirmation of EV markers on CVL and DC EVs. Shown are averages of tetraspanin-positive particles bound to anti-CD63 and anti-CD81 antibodies and detected by label-free imaging.

### Study Approvals

All animal studies were approved by the Johns Hopkins University Institutional Animal Care and Use Committee (IACUC) and conducted in accordance with the Weatherall Report, the Guide for the Care and Use of Laboratory Animals, and the USDA Animal Welfare Act.

### Sample Processing

Sample processing began within a maximum of 60 minutes of collection and utilized serial centrifugation steps to enrich EVs as described previously ^30^, based on a standard EV isolation protocol ^43^. Specifically, fluids were centrifuged: (1) 1,000 × *g* for 15mins at 4°C in a tabletop centrifuge; (2) 10,000 × *g* for 20 mins at 4°C; and (3) 110,000 × *g* for 2 hours at 4°C with a Sorvall Discovery SE ultracentrifuge (Thermo Fisher Scientific) with an AH-650 rotor (k factor: 53.0) (Supplemental Figure 1B). Following each centrifugation step, most supernatant was removed, taking care not to disturb the pellet. After each step, supernatant was set aside for nanoparticle tracking analysis (NTA; 200 µL), and RNA isolation (200 µL) following the second and third steps. The pellet was resuspended in 400 µL of PBS after each centrifugation step. After the final step, the remaining ultracentrifuged supernatant was concentrated to approximately 220 µL using Amicon Ultra-2 10 kDa molecular weight cutoff filters (Merck KGaA, Darmstadt, Germany. Cat #: UFC201024). 200 µL of the concentrate was used for RNA isolation and the remainder was retained for NTA. All samples reserved for RNA isolation were mixed with 62.6 µL of RNA isolation buffer (Exiqon, Vedbaek, Denmark. Cat #: 300112. Lot #: 593-84-9n) containing three micrograms of glycogen and 5 pg of synthetic cel-miR-39 as previously described ^44^. Processed samples were analyzed immediately or frozen at −80°C until further use.

For plasma, whole blood was centrifuged at 800 × *g* for 10 mins at 25°C. Supernatant was centrifuged twice at 2,500 × *g* for 10 mins at 25°C. The resulting platelet-poor plasma was aliquoted and frozen at −80°C.

### Hormone Analysis

Levels of progesterone (P4) and estradiol-17b (E2) were measured in plasma samples shipped overnight on dry ice to the Endocrine Technology and Support Core Lab at the Oregon National Primate Research Center, Oregon Health and Science University.

### Nanoparticle Tracking Analysis

Extracellular particle concentration was determined using a NanoSight NS500 NTA system (Malvern, Worcestershire, UK). Cervicovaginal lavage samples were diluted as needed and specified in Supplemental Table 2 to ensure optimal NTA analysis. At least five 20-second videos were recorded for each sample at a camera setting of 12. Data were analyzed at a detection threshold of two using NanoSight software version 3.0.

### Western Blot

Western blot was used to detect the presence of EV protein markers and the absence of nucleoporin (nuclear marker) in CVL and enriched CVL EVs. 20 µL of samples from each fraction were lysed with 5 µL 1:1 mixture of RIPA buffer (Cell Signaling Technology, Danvers, MA. Cat #: 9806S) and protease inhibitor (Santa Cruz Biotechnology, Dallas, TX. Cat #: sc29131). 8 µL of Laemmli 4X sample buffer (BioRad, Hercules, CA. Cat #:161-0747 Lot #: 64077737) was added per sample, and 30 µL of each was loaded into a Criterion TGX 4-15% gel (BioRad, Hercules, CA. Cat #: 5678084 Lot #: 64301319) after 5 mins of 95 °C incubation. The gel was electrophoresed by application of 100V for 100 mins. The proteins were then transferred to a PVDF membrane (BioRad, Hercules, CA. Cat #: 1620177, Lot #:31689A12.), which was blocked with 5% milk (BioRad, Hercules, CA. Cat #: 1706404. Lot #: 64047053) in PBS+0.1%Tween®20 (Sigma-Aldrich, St. Louis, MO Cat #: 274348 Lot #: MKBF5463V) for 1 hour. The membrane was subsequently incubated with mouse anti-human CD63 (BD Biosciences, San Jose, CA Cat #: 556019 Lot #: 6355939) and mouse monoclonal IgG_2b CD81 (Santa Cruz Biotechnology, Dallas, TX Cat #: 166029 Lot #: L1015) primary antibodies, at a concentration of 0.5 µg/mL overnight. After washing the membrane, it was incubated with a goat anti-mouse IgG-HRP secondary antibody (Santa Cruz Biotechnology, Dallas, TX Cat #: sc-2005 Lot #: B1616) at a 1:5,000 dilution for 1 h. The membrane was then incubated with a 1:1 mixture of SuperSignal West Pico Stable Peroxide solution and Luminol Enhancer solution (Thermo Scientific, Rockford, IL Cat #: 34080 Lot #: SD246944) for 5 min. The membrane was visualized on Azure 600 imaging system (Azure Biosystems, Dublin, CA). The second blot was done in a reducing environment using 10mM DTT (Promega, Madison, WI Cat #: P1171 Lot #: 0000198991). Same procedures were followed with rabbit anti-human TSG101 (Cat #: ab125011 Lot #:GR180132-14), rabbit polyclonal anti-nucleoporin (Abcam, Cambridge, MA Cat #: ab96134 Lot #: GR22167-18) primary antibodies. Subsequent incubation with goat anti-rabbit IgG-HRP secondary antibody (Abcam, Cambridge, MA Cat #: sc-2204 Lot #: B2216). All antibodies were used at the same concentration as the first blot. Membrane was visualized on the Azure imaging system.

### Single particle interferometric reflectance imaging

Both CVL-derived and dendritic cell LK23-derived EVs were diluted 1:1000 and incubated on ExoView (NanoView Biosciences, Brighton, MA) chips that were printed with anti-CD63 (BD Biosciences, Bedford, MA. Cat#: 556019) and anti-CD81 (BD Biosciences, Bedford, MA. Cat #: 555675) antibodies. After incubation for 16 hours, chips were washed per manufacturer’s protocol and imaged in the ExoView scanner by interferometric reflectance imaging.

### Electron Microscopy

Gold grids were floated on 2% paraformaldehyde-fixed CVL–derived samples for two minutes, then negatively stained with uranyl acetate for 22 seconds. Grids were observed with a Hitachi 7600 transmission electron microscope in the Johns Hopkins Institute for Basic Biomedical Sciences Microscope Facility.

### Total RNA Isolation and Quality Control

RNA isolation work flow is shown in Supplemental Figure 1C. RNA lysis buffer was added into each sample as described above prior to freezing (−80 °C). Total RNA was isolated from thawed samples using the miRCURY RNA Isolation Kit-Biofluids (Exiqon, Vedbaek, Denmark. Cat #: 300112. Lot #: 593-84-9n) per manufacturer’s protocol with minor modifications as previously described^44^. Total RNA was eluted with 50 µL RNase-free water and stored at −80°C. As quality control, expression levels of several small RNAs including snRNA U6, miR-16-5p, miR-223-3p, and the spiked-in synthetic cel-miR-39 were assessed by TaqMan miRNA assays (Applied Biosystems/ Life Technologies, Carlsbad, California, USA) ^45^.

### miRNA Profiling by TaqMan Low-Density Array

A custom 48-feature TaqMan low-density array (TLDA) was ordered from Thermo Fisher, with features chosen based on results of a human CVL pilot study (GVH and KWW, unpublished data). Stem-loop primer reverse transcription and pre-amplification steps were conducted using the manufacturer’s reagents as previously described ^46^ but with 14 cycles of pre-amplification. Real time quantitative PCR was performed with a QuantStudio 12K instrument (Johns Hopkins University DNA Analysis Facility). Data were collected using SDS software and Cq values extracted with Expression Suite v1.0.4 (Thermo Fisher Scientific, Waltham, MA USA). Raw Cq values were adjusted by a factor determined from the geometric mean of 10 relatively invariant miRNAs. The selection process for these invariant miRNAs was to 1) rank miRNAs by coefficient of variation; 2) remove miRNAs with high average Cq (>30), non-miRNAs, and those with low amplification score; 3) select the lowest-CV member of miRNA families (e.g., the 17/92 clusters); and 4) pick the top 10 remaining candidates by CV: let-7b-5p, -miR-21-5p, -27a-3p, -28-3p, -29a-3p, -30b-5p, -92a-3p, -197-3p, -200c-3p, and -320a-3p.

### Individual RT-qPCR Assays

Individual TaqMan miRNA qPCR assays were performed as previously described ^46^ on all UC pellet samples from all animals across all weeks for miRs-19a-3p (Thermo Fisher Assay ID #000395), -186-5p (Thermo Fisher Assay ID #002285), -451a-5p (Thermo Fisher Assay ID #001105), -200c-3p (Thermo Fisher Assay ID #002300), -222-3p (Thermo Fisher Assay ID #002276), -193b-3p (Thermo Fisher Assay ID #002367), -181a-5p (Thermo Fisher Assay ID #000480), -223a-3p (Thermo Fisher Assay ID #002295), -16-5p (Thermo Fisher Assay ID #000391), -106a-5p (Thermo Fisher Assay ID #002169), and -125b-5p (Thermo Fisher Assay ID #00449). We also measured miR-375-3p (Thermo Fisher Assay ID #00564), which was not included on the array. Data were adjusted to Cqs of miR-16-5p.

### Blood Cell Isolation and Monocyte-Derived Macrophage Culture

Total PBMCs were obtained from freshly drawn blood from human donors under a Johns Hopkins University School of Medicine IRB-approved protocol (JHU IRB #CR00011400). Blood was mixed with 10% Acid Citrate Dextrose (ACD) (Sigma Aldrich, St. Louis, MO Cat #: C3821 Lot #: SLBQ6570V) with gentle mixing by inversion. Within 15 minutes of draw, blood was diluted with equal volume of PBS+ 2% FBS, gently layered onto room temperature Ficoll (Biosciences AB, Uppsala, Sweden Cat #:17-1440-03 Lot #: 10253776) in Sepmate-50 tubes (STEMCELL Technologies, Vancouver, BC, Canada Cat #: 15450 Lot #: 06102016) and centrifuged for 10 minutes at 1200 × *g*. Plasma and PBMC fractions were removed, washed in PBS+ 2% FBS, and pelleted at 300 × *g* for 8 minutes. Pellets from 5 tubes were combined by resuspension in 10 mL RBC lysis buffer (4.15 g NH_4_Cl, 0.5 g KHCO_3_, 0.15 g EDTA in 450 mL H_2_O; pH adjusted to 7.2–7.3; volume adjusted to 500 mL and filter-sterilized); total volume was brought to 40 mL with RBC lysis buffer. After incubation at 37 °C for 5 mins, the suspension was centrifuged at 400 × *g* for 6 mins at room temperature. The cell pellet was resuspended in Macrophage Differentiation Medium with 20% FBS (MDM20) to a final concentration of 2×10^6^ cells/mL. PBMCs were plated at 4×10^6^ cells per well in 12-well plates and cultured in MDM20 for 7 days. One half of the total volume of medium was replaced on day 3. On day 7, cells were washed 3 times with PBS to remove non-adherent cells. The medium was replaced with Macrophage Differentiation Medium with 10% serum (MDM10) and cultured overnight prior to transfection.

### miRNA Mimic Transfection

Differentiated macrophages were transfected with 50 nM miRNA-186-5p (Qiagen, Foster City, CA. Cat #: MSY0000456 Lot #: 286688176) using Lipofectamine 2000 (Invitrogen/Life Technologies, Carlsbad, CA Cat #: 11668-019 Lot #:1467572) diluted in OptiMEM Reduced Serum Medium (Gibco, Grand Island, NY Cat #: 31985-070 Lot #: 1762285). Controls included mock transfections and transfection of 50 nM double-stranded siRNA oligo labeled with Alexa Fluor 555 (Invitrogen, Fredrick, MD Cat #: 14750-100 Lot #: 1863892). Plates were incubated for 6 hours at 37 °C. After incubation, successful transfection was confirmed by examining uptake of labeled siRNA with an Eclipse TE200 inverted microscope (Nikon Instruments, Melville, NY). Transfection medium was removed. The plates were washed with PBS and refed with 2 mL fresh MDM10 medium.

### HIV Infection

HIV-1 BaL stocks were generated from infected PM1 T-lymphocytic cells and stored at −80 °C. 24 hours after mimic or mock transfections, macrophages were infected with HIV BaL and incubated overnight (stock, 80 µg p24/mL, diluted to 200 ng p24/mL). At days 3, 6, and 9 post-infection, 500 µL supernatant was collected for p24 release assays and cells were lysed with 600 µL mirVana lysis buffer for subsequent RNA isolation and analysis.

### HIV p24 Antigen ELISA

Supernatant samples were lysed with Triton-X (Perkin Elmer, Waltham, MA Cat #: NEK050B001KT Lot #: 990-17041) at a final concentration of 1%. The DuPont HIV-1 p24 Core Profile ELISA kit (Perkin Elmer, Waltham, MA Cat #: NEK050B001KT Lot #: 990-17041) was used per manufacturer’s instructions to measure p24 concentration based on the provided standard.

### Total RNA Isolation

Total RNA was isolated using the mirVana miRNA Isolation Kit per manufacturer’s protocol (Ambion, Vilnius, Lithuania. Cat #: AM1560 Lot #: 1211082). Note that this procedure yields total RNA, not just small RNAs. After elution with 100 µL RNase-free water, nucleic acid concentration was measured using a NanoDrop 1000 spectrophotometer (Thermo Fisher Scientific, Wilmington, DE). RNA isolates were stored at −80 °C.

### HIV Gag RNA RT-qPCR

Real-time one-step reverse transcription quantitative PCR was performed with the QuantiTect Virus Kit (Qiagen, Foster City, CA Cat #:211011 Lot #: 154030803). Each 25 µl reaction mixture contained 15 µl of master mix containing HIV-1 RNA standard, 100 µM of FAM dye and IBFQ quencher labeled Gag probe (5’ ATT ATC AGA AGG AGC CAC CCC ACA AGA 3’), 600 nM each of Gag1 forward primer (5’TCA GCC CAG AAG TAA TAC CCA TGT 3’) and Gag2 reverse primer (5’ CAC TGT GTT TAG CAT GGT GTT T 3’), nuclease-free water, and QuantiTect Virus RT mix, and 10 µL serial-diluted standard or template RNA. No-template control and no reverse transcriptase controls were included. Linear standard curve was generated by plotting the log copy number versus the quantification cycle (Cq) value. Log-transformed Gag copy number was calculated based on the standard curve.

### Data analysis

Data processing and analysis were conducted using tools from Microsoft Excel (geometric mean normalization), Apple Numbers, GraphPad Prism, the MultiExperiment Viewer, and R/BioConductor packages including pheatmap (http://CRAN.R-project.org/package=pheatmap; quantile normalization, Euclidean distance, self-organizing maps, self-organizing tree algorithms, k-means clustering). Figures and tables were prepared using R Studio, Microsoft Excel and Word, Apple Numbers and Keynote, GraphPad Prism, and Adobe Photoshop.

### Data Availability and Rigor and Reproducibility

Array data have been deposited with the Gene Expression Omnibus (GEO) ^47^ as GSE107856. Data in other formats are available upon request. To the extent that sample quantities would allow, the MISEV recommendations for EV studies were followed^24^, and the EV experiments have been registered with the EV-TRACK knowledgebase, ^48^ with preliminary EV-TRACK code XL5296IL.

## 3. Results

### 3.1. Abnormal menstrual cycle of SIV-infected macaques and ovulation-associated changes in CVL EV-enriched particles

Plasma and CVL were collected from two control and four SIV-infected macaques over the course of five weeks (Supplemental Figure 1). Abnormal cycling was observed for infected subjects (K. Mulka, et al, unpublished data). By nanoparticle tracking analysis, CVL EV concentration in control animals increased during ovulation (Figure 1A). Transmission electron microscopy was performed for representative fractions of CVL, revealing bacteria and large particles in the 10,000 × g pellet (Figure 1B). The 100,000 x g pellet included apparent EVs up to 200 nm in diameter (Figure 1C). EV markers (shown: CD63, CD81, and TSG101) were confirmed by Western blot (Figure 1D). The nuclear marker nucleoporin was detected only in tissue samples (Figure 1D). The relative EV tetraspanin profiles of both CVL and control EV samples were corroborated with single particle interferometric reflectance imaging: CVL EVs had a higher CD63 expression and dendritic cell EVs had higher CD81 expression.

### 3.2. TLDA reveals an extracellular miRNA profile of the cervicovaginal compartment

Based upon preliminary findings from a study of human CVL (Hancock and Witwer, unpublished data), we designed a custom TaqMan low-density array (TLDA) to measure 47 miRNAs expected to be present in CVL, along with the snRNA U6. CVL from all subjects and at all time points was fractionated by stepped centrifugation to yield a 10,000 x g pellet (10K pellet), a 100,000 x g pellet (UC pellet), and 100,000 x g supernatant (UC supernatant). Total RNA from all fractions was profiled by TLDA. Raw (Supplemental Figure S2A), quantile normalized (Supplemental Figure S2B), and geometric mean-adjusted Cq values (Supplemental Figure S2C) were subjected to unsupervised hierarchical clustering. This clustering did not reveal broad miRNA profile differences associated with sample collection time, menstruation, or SIV infection.

### 3.3. Distribution of miRNAs across CVL fractions

Across the three examined CVL fractions (p10, p100, S100), the ten most abundant miRNAs (lowest Cq values) were miRs-223-3p, -203a-3p, -24-3p, -150-5p, -21-5p, -146a-5p, -92a-3p, -222-3p, - 17-5p, and -106a-5p. The average normalized Cq value for each miRNA was greater (i.e., lower abundance) in the p100 than the s100 fraction (Figure 2A and inset), and indeed in p10 and p100 combined (Figure 3B), suggesting that most miRNA in CVL, as reported for various other body fluids, is found outside the EV-enriched fractions. Considering all fractions, the differences between the EV-enriched and EV-depleted fractions were significant even after Bonferroni correction for all features except U6. On average, the s100 fraction contained 86.5% of the total miRNA from these three fractions. In the p10 fraction, the average miRNA was detected at 10.5% its level in the s100 fraction (SD=5.7%). miR-34a-5p had the lowest (5.9%) and miR-28-3p the highest (33.7%) abundance compared with s100. In the p100 fraction, miRNAs were on average 5.6% (SD=2.4%) as abundant as in s100. The least represented in p100 was miR-27a-3p (2.3%), and the best represented was again miR-28-3p (13.4%). Together, the content of the EV-enriched fractions (p10 and p100) as a percentage of the total is shown in Figure 2B for individual miRNAs. miRNA rank was significantly correlated across fractions, despite minor differences in order (Figure 2C).

**Figure 2.**
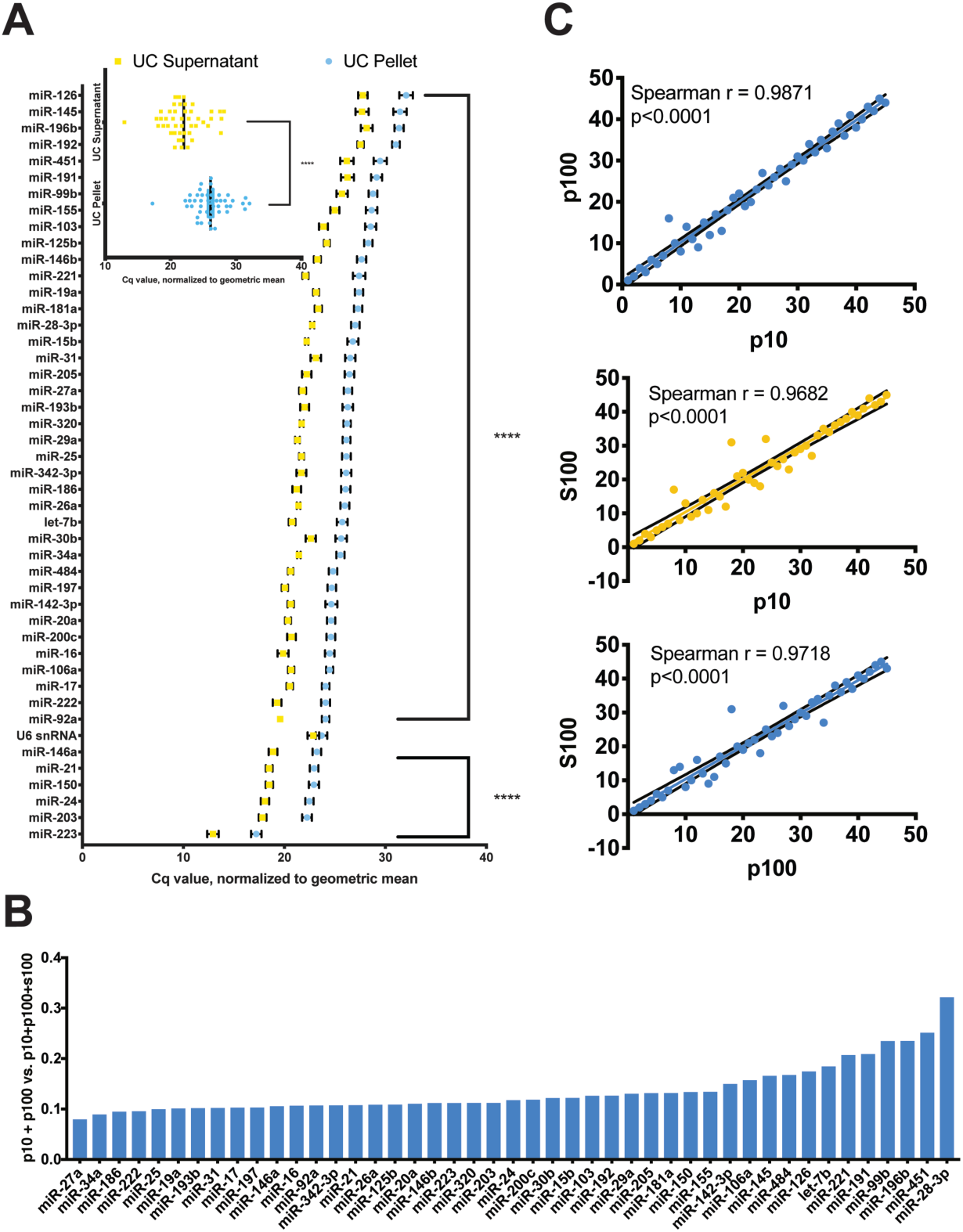
Relative abundance of miRNAs in different CVL fractions. **A)** Abundant miRNAs in descending order based on Cq values normalized to the geometric mean for each sample. Inset: average of all miRNAs in UC pellet and UC supernatant. Error bars: SEM. **B)** miRNA expression in EV-enriched fractions (p10, p100) as a percentage of total estimated expression (p10+p100+S100 by Cq) in ascending order, from miR-27a-3p (7.9%) to miR-28-3p (32.0%). **C)** miRNAs in each fraction (10,000× g pellet=p10, 110,000 × g pellet=p100, 110,000× g supernatant=S100, and) are significantly correlated (p<0.0001, Spearman).

**Figure 3.**
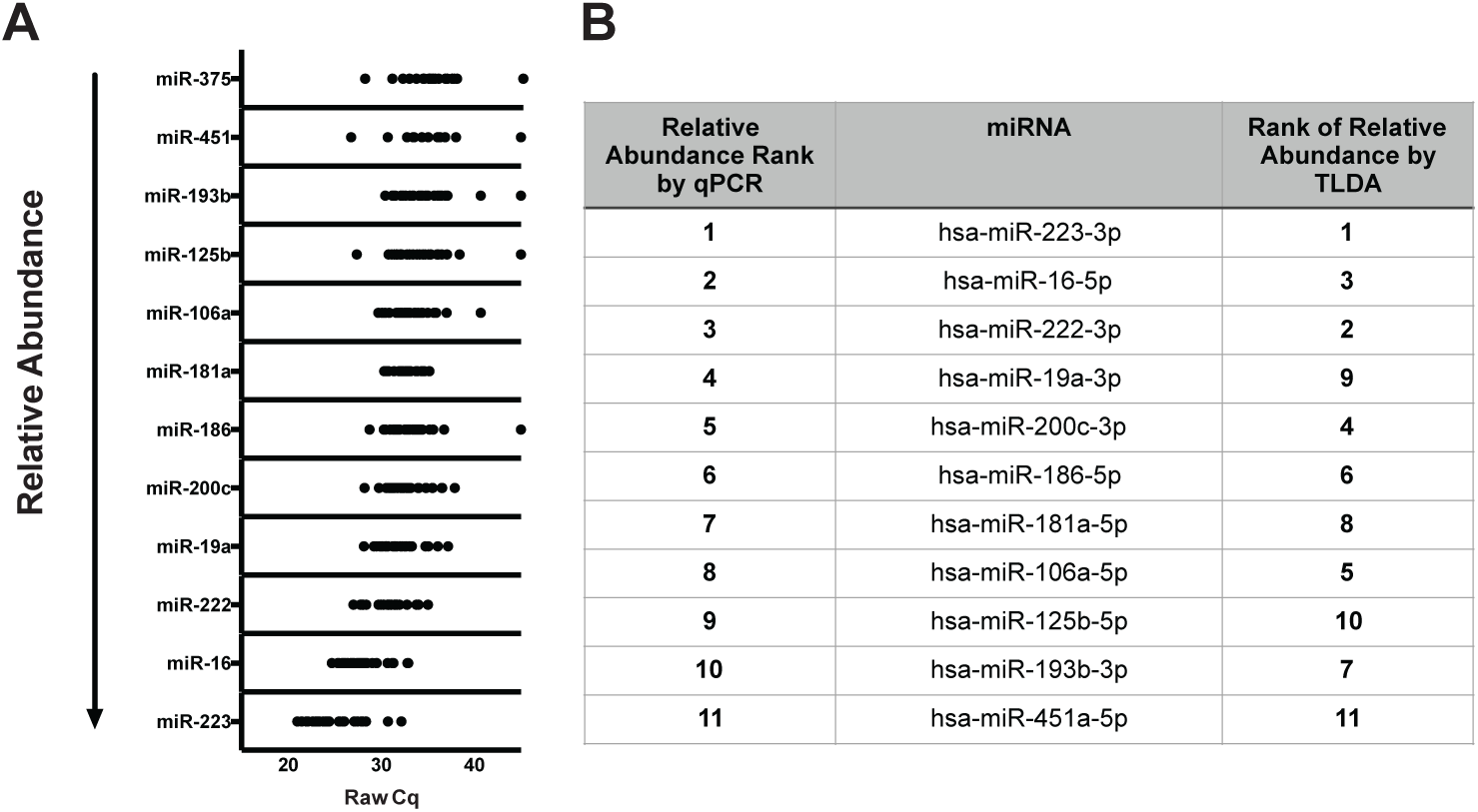
miRNA qPCR validation. **A)** qPCR validation for UC pellet samples, all subjects and time points (individual dots). **B)** Ranks of abundant miRNAs by qPCR and TLDA.

### 3.4. qPCR Validation

Individual stem loop RT/hydrolysis probe qPCR assays were used to verify TLDA results for eleven selected miRNAs plus miR-375-3p (not included on the array), which was also measured because of a reported association with goblet cells ^49^. Some miRNAs were chosen due to high expression levels. miR-181a-5p was measured due to its association with endometrial cells ^50,51^. miR-125b-5p has been reported as a diagnostic marker of endometriosis ^52^. Other miRNAs (miRs-186-5p, -451a-5p, -200c-3p, -222-3p, -193b-3p) were selected based on our previous experience and results from other studies evaluating miRNAs in the context of HIV-1 and SIV infections. Results of qPCR assays, adjusted by miR-16-5p for each sample (since we found relatively low qPCR variation of miR-16-5p, a commonly used normalizer ^53^), are shown in Figure 3A. Figure 3B compares miRNA ranks (1-11) by TLDA and individual qPCR, which are generally in concordance. Note that expression of red blood cell miRNA miR-451a-5p was low, suggesting minimal contamination from blood for most samples.

### 3.5. miRNA association with retroviral infection status

An association of miRNA abundance with infection status could yield novel biomarkers as well as clues to roles of miRNA in modulating infection. However, the small number of subjects in our study was a challenge. Nevertheless, by considering all subjects and time points together for both infected and uninfected subjects, microarray data suggested a slightly reduced abundance of miRs-186-5p, -222-3p, and -200c-3p in infected samples (Figure 4A) based on statistical analysis of Δ*Cq* values, while qPCR revealed differential abundance of miRs-186-5p and -125b-5p (Figure 4B). miR-186-5p was thus identified by both techniques as potentially associated with retroviral infection.

**Figure 4.**
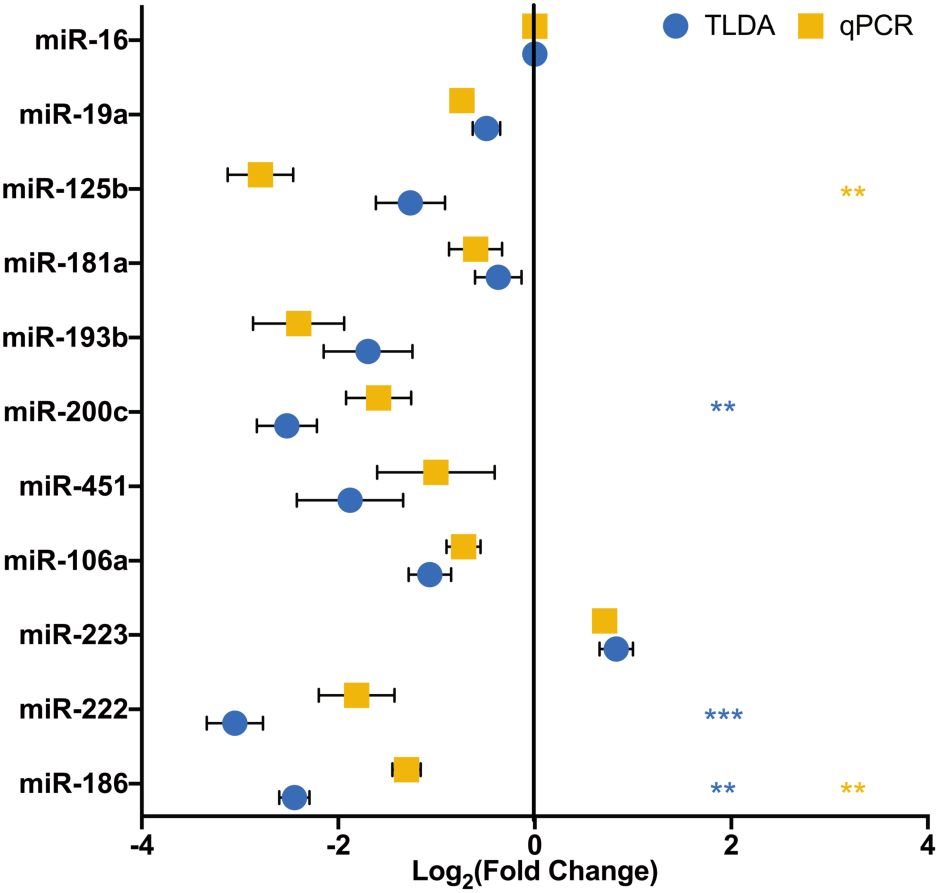
miR-186-5p downregulation: SIV. miR-186-5p fold change was determined using ΔΔ*Ct* method using miR-16 and uninfected animals as controls. *Log2(fold change)* for both TLDA and qPCR analyses was plotted for 11 selected validation miRNAs. Statistical analyses were performed on Δ*Ct* values. For TLDA, miRs-186, −222, and −200c were significantly less abundant in the CVL p100 fraction of infected subjects (Mean ± SEM, Multiple t test, Bonferroni-Dunn Correction), ** p < 0.01, *** p<0.001. For qPCR analysis, miRs-186 and −125b were significantly less abundant (Multiple t test, Bonferroni-Dunn Correction), ** p < 0.01, *** p<0.001

### 3.6. miR-186-5p transfection has minimal effects on cellular HIV RNA abundance but reduces p24 release from monocyte-derived macrophages

To assess a possible influence of miR-186-5p (“miR-186”) on retroviral replication, we introduced double-stranded miR-186-5p mimic or control RNA into monocyte-derived macrophages derived from three donors 24 hours before infecting the cells or not with HIV. At days three and six post-infection, we quantitated full-length HIV-1 transcript using a gag qPCR with standard curve. In cells from only one of three donors were fewer HIV-1 copies associated with miR-186-5p mimic transfection (Figure 5). Overall, there was no statistically significant difference in HIV RNA between the conditions.

**Figure 5.**
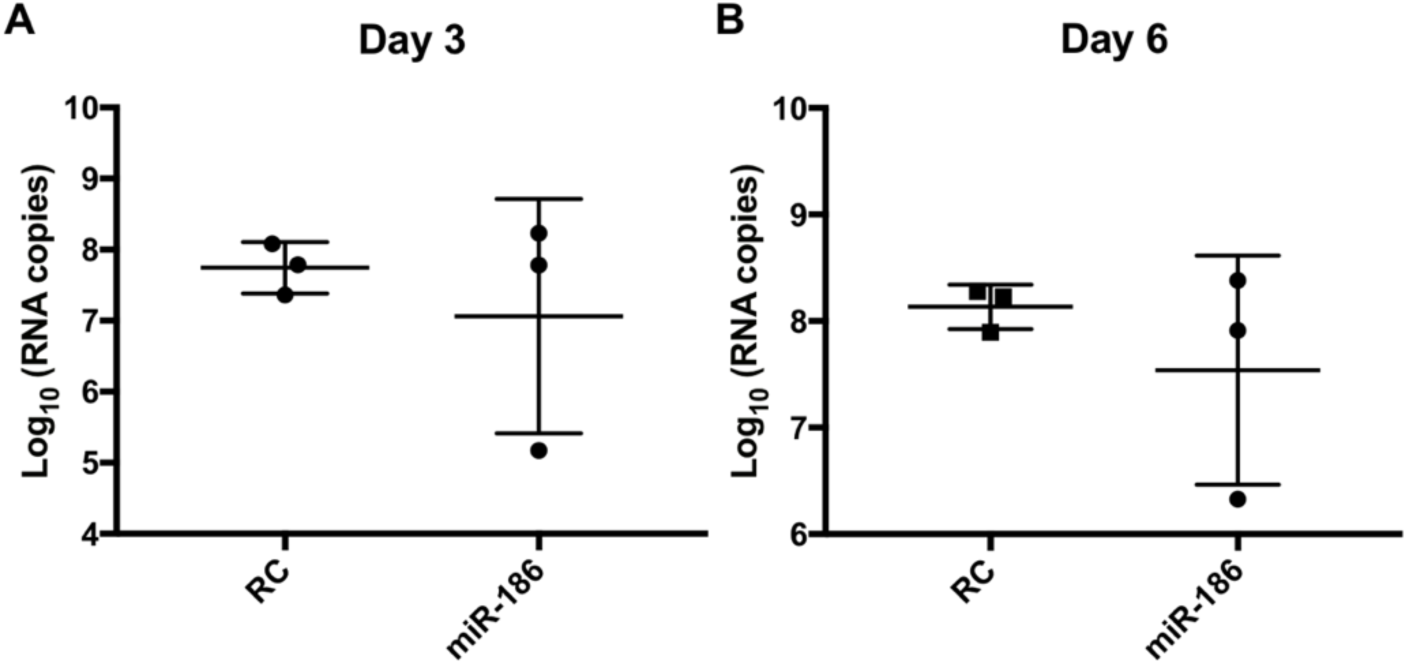
miRNA-186-5p mimic transfection inconsistently suppresses HIV-1 gag mRNA production. Apparent downregulation of gag mRNA (qPCR assay with standard curve) was observed in miR-186-transfected monocyte-derived macrophages from only 1 of 3 donors compared with control RNA-transfected cells (RC). Overall, results were insignificant by t-test, p>0.1, with multiple replicates of cells from 3 human donors.

However, at the same time points and also out to nine days post-infection, a different result was seen for capsid p24 release into the supernatant. For infected but untransfected cells, measurable p24 was observed by 3 dpi, and p24 counts increased by two-fold or more by 9 dpi (Figure 6A) for multiple replicate experiments with cells from three donors. Compared with infected, untreated controls, mock-transfected cells (not shown), and cells transfected with a negative control RNA (labeled with a fluorophore to assess transfection efficiency), miR-186-5p transfection was associated with a significant decline of released p24 at all time points (ANOVA with Bonferroni correction) (Figure 6B-D). The negative control condition showed a suppressive trend that reached nominal significance at 9 dpi. However, miR-186-associated suppression was significantly greater at all time points.

**Figure 6.**
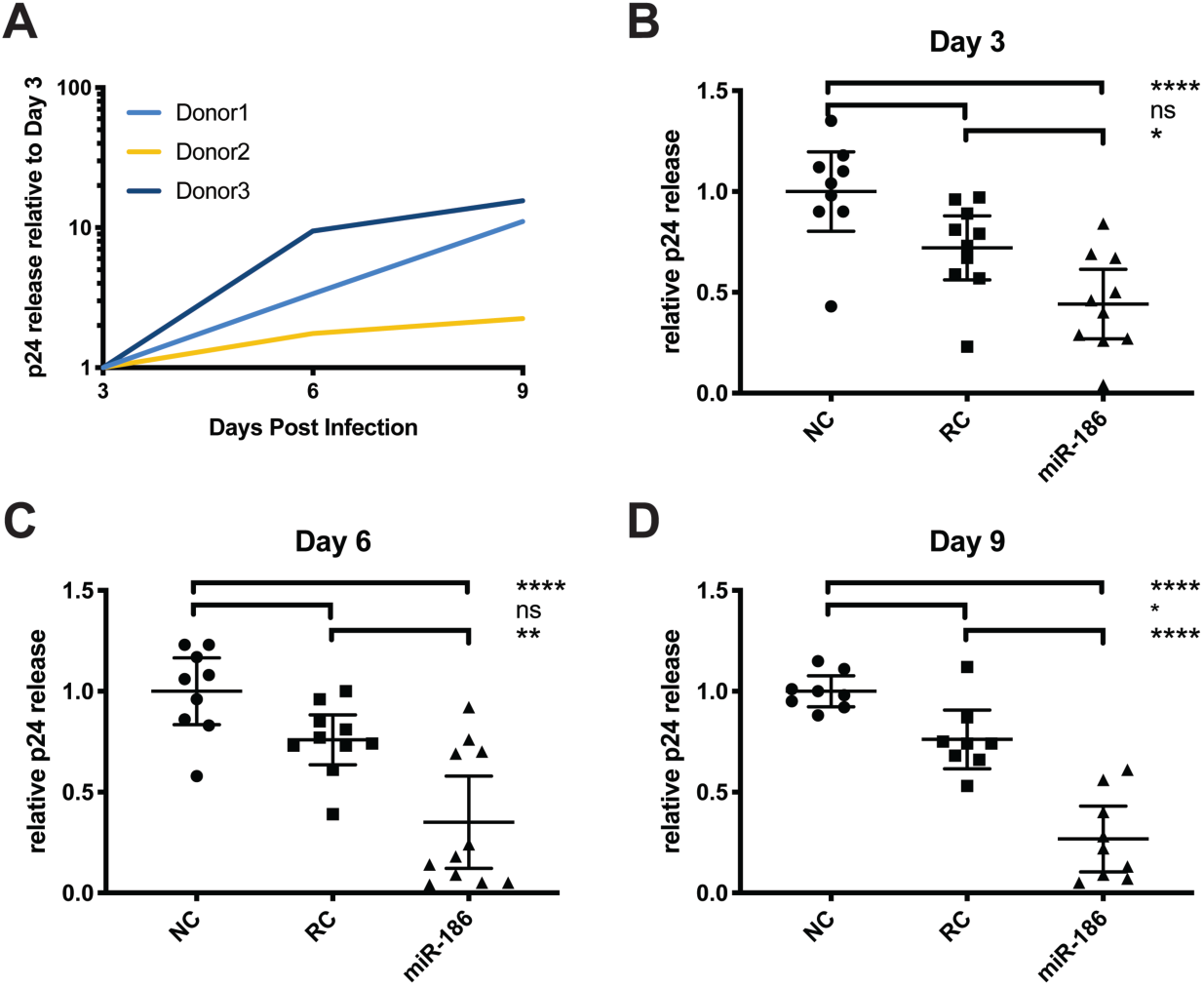
miRNA-186-5p inhibits p24 release. Monocyte-derived macrophages from human donors were infected with HIV-1 BaL. **A)** p24 production increased >2 fold for all donors from 3 to 9 days post-infection (dpi), untreated cells. **B-D)** Transfection of miR-186-5p mimic was associated with a decrease of p24 release compared with untransfected controls (NC) and control RNA mimic-transfected controls (RC) at the indicated time points; ns=not significant, * p<0.05, ** p<0.01, **** p<0.0001 (ANOVA followed by Bonferroni correction for multiple tests). Results were from 8 to 11 replicate experiments with cells from 3 human donors.

### 3.7 p24 inhibition by miR-186-5p is correlated with transfection efficiency

Despite the statistical significance of miR-186-5p-associated p24 inhibition, substantial variability was observed, including between donors/experiments; we therefore hypothesized that either donor- or experiment-specific factors were responsible for the variability. The transfection experiments were repeated using macrophages from five additional donors (labeled 1-5). While significant but variable inhibition of p24 release after miR-186-5p transfection was observed for three donors (1, 2, and 5), little or no inhibition was seen for donors 3 and 4 (Figure 7).

**Figure 7.**
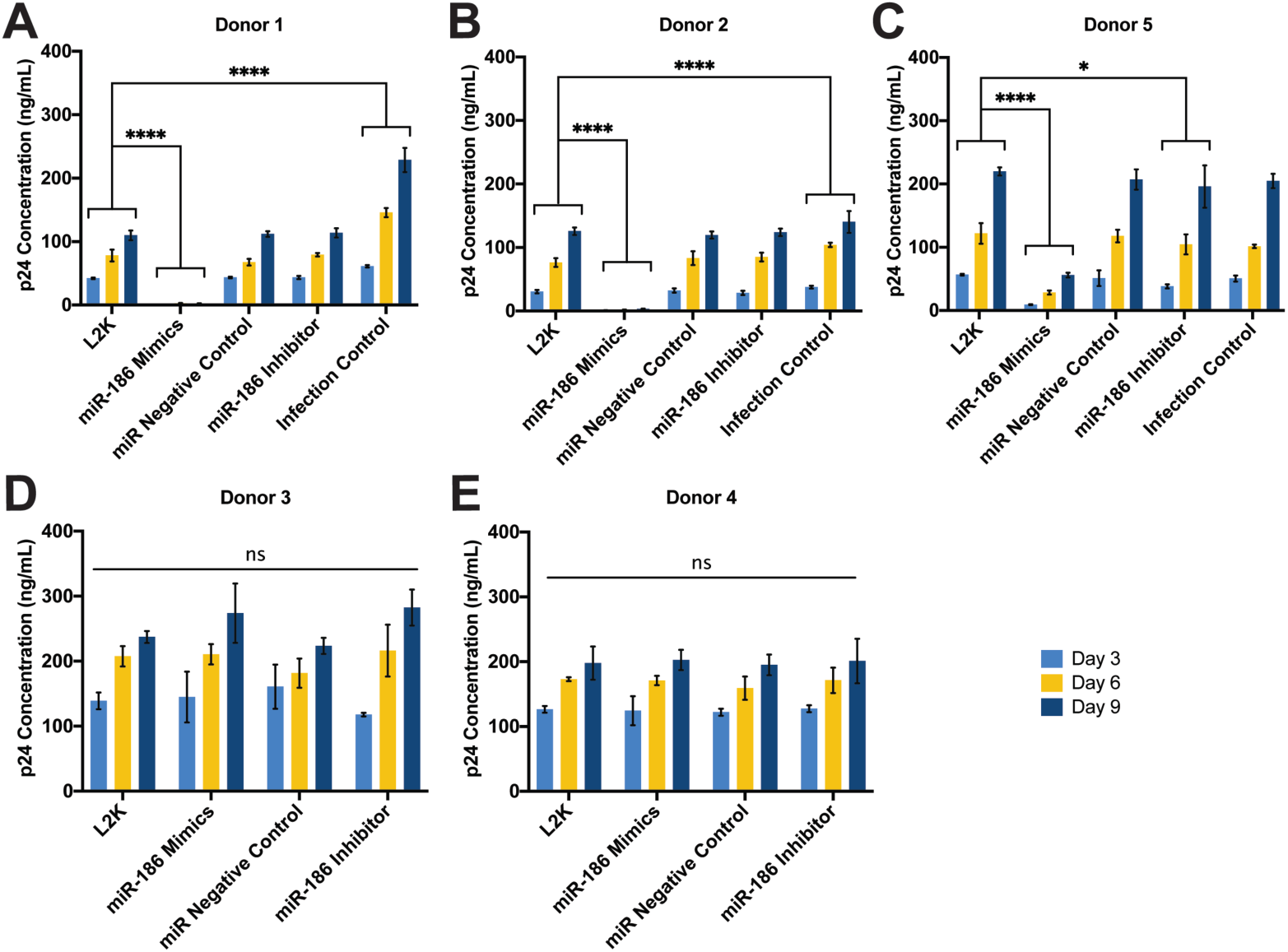
miRNA-186-5p inhibits p24 release in a donor-specific manner. Monocyte-derived macrophages from human donors were infected with HIV-1 BaL. **A-C)** Compared with mock transfected controls, transfection of miR-186 mimic was associated with a significant decrease of p24 production from 3 to 9 days post-infection (dpi) in donors 1, 2, and 5. **D-E)** For donors 3 and 4, transfection of miR-186 mimic was ineffective in inhibiting p24 release compared with mock transfected controls; ns=not significant, * p<0.05, ** p<0.01, **** p<0.0001 (Two-way ANOVA followed by Bonferroni correction for multiple tests).

One experimental variable that could affect the degree of inhibition is the efficiency with which the miRNA mimic is delivered into the cells. Since this variable was not assessed in our previous experiments, we measured it for the five new experiments. Despite using the same nominal concentrations of miRNA mimics for our experiments, a nearly 100-fold range of miR-186-5p concentration was observed between the lowest- and highest-efficiency transfections (Figure 9A), which increased miR-186-5p levels from around 5-fold to nearly 500-fold, respectively. Strikingly, the miR-186-5p level was inversely correlated with released p24 across these five donors. It should be noted that miR-186-5p antisense inhibitors were also introduced in these experiments. While they did not significantly increase HIV p24 release (Figure 7), they also did not achieve a consistent knockdown of native miR-186-5p (Figure 8A).

**Figure 8.**
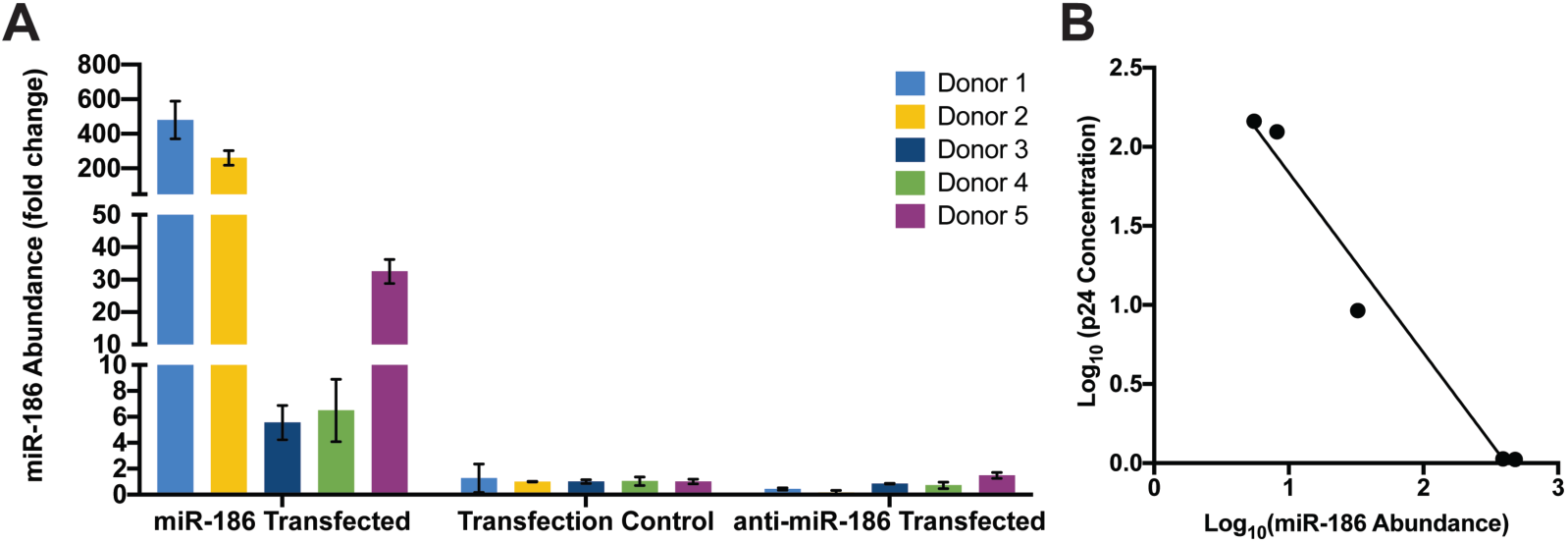
miR-186-5p abundance post-transfection and correlation with p24 release. **A)** Abundance of miR-186-5p in macrophages post-transfection, as assessed by qPCR and compared (fold change) with the average of control macrophages. **B)** Correlation of macrophage miR-186-5p and p24 concentration released in supernatant three days post-infection. p(two-tailed)=0.0019 (Correlation), *R*^*2*^ = 0.9731(Linear regression).

## 4. Discussion

Cervicovaginal lavage EVs and exRNPs, like EVs in the uterus ^54,55^, may offer information about the health of the reproductive tract and may also facilitate or block transmission of infectious agents. Proteomic analyses of human ^56^ and rhesus macaque ^57^ CVL have suggested a core proteome and a highly variable proteome that responds to changes in pregnancy status, menstruation, infection, and other stressors. However, exRNA and extracellular vesicle profiles are less understood in this compartment. Thus, one major finding of this study is a partial profile of miRNAs of EV-enriched and -depleted fractions of CVL fluid of primates. We report that EVs can be liberated from vaginal secretions by lavage, and that these EVs can be concentrated using a standard stepped centrifugation procedure, with enrichment of positive (membrane-associated) markers while a cellular negative control was not detected.

Both EV-replete and EV-depleted fractions of CVL contained abundant miRNA. As reported for other biological fluids ^37,58^, miRNA concentration was highest in the EV-depleted CVL fractions, not in EV-enriched ultracentrifuged pellets, consistent with packaging of most extracellular miRNA into exRNPs; the function, if any, of extracellular miRNAs in the cervicovaginal tract of healthy individuals remains to be determined. We observed minimal differences in extracellular miRNA profiles between SIV-infected and uninfected subjects or, surprisingly, even during the menstrual cycle, suggesting a certain stability of extracellular miRNA in the compartment. Correlation of miRNA concentrations in EV-depleted and -replete fractions was also apparent. Based on relative abundance compared with miRNAs of other cellular/tissue origins (e.g., heart and lung specific miR-126, kidney-specific miR-196b, and liver-specific miR-192) ^59,60^, miRNAs in EVs and exRNPs of CVL are likely derived from epithelial cells (including goblet cells), and cells of the immune system (as suggested, e.g., by myeloid-enriched miR-223 and lymphocyte-enriched miR-150)^61^. Of the most abundant miRNAs we identified, some have been ascribed tumor-suppressive roles in cancers^62–68^. Also, miR-223 and miR-150 have been described as “anti-HIV” miRNAs ^69^ among a variety of reported antiretroviral small RNAs (sRNAs), both host and viral ^70–75^. Given their relative abundance in the vaginal tract, a common site for HIV infection, these miRNAs may contribute to antiviral defenses.

Along these lines, a second major finding of this study is a possible role for miR-186-5p in antiretroviral defense, bolstered by the observation that exogenous miR-186-5p transfection efficiency correlates inversely with HIV p24 release. Previous publications have identified protein constituents in the cervicovaginal lavage with anti-HIV efficacies (for example ^7,14,22^). Our identification of miRNA as a potential anti-HIV agent adds an element of complexity to the picture of tissue-specific antiretroviral defense. In contrast with an early report of direct binding of host miRNAs to retroviral transcripts and subsequent suppression ^69^, it now appears that this mechanism of suppression may be relatively uncommon ^76^. Anti-HIV miRNAs may be more likely to exert effects through control of host genes instead (e.g., ^77^). Our data also support the conclusion that reduction of HIV RNA levels is not the main mechanism for miR-186-mediated suppression of HIV release.

How, then, might miR-186-5p, whether endogenous or exogenous (therapeutically introduced) contribute to antiretroviral effects? Combining several miRNA target prediction, validation, and enrichment analysis approaches ^78–84^, we noticed a few putative miR-186-5p targets and related pathways that may merit follow-up. One target of miR-186-5p that was validated experimentally by multiple methods is FOXO1 ^85^, an important contributor to apoptosis but also immunoregulation via IFNγ pathways. Another prominent validated target, P2X7R ^86^, is involved in membrane budding, T-cell-mediated cytotoxicity, cellular response to extracellular stimuli and T-cell homeostasis/proliferation. There is also evidence that miR-186-5p targets the HIV co-receptor CXCR4 ^87^. Pathway enrichment analyses ^83,84^ suggest that miR-186-5p targets participate significantly in infection-related networks, including prion diseases, viral carcinogenesis, and responses to measles and herpes simplex virus infections. Although miRNA target prediction algorithms are imperfect, and validation efforts are of varying quality ^88,89^, these findings may shed some light on how miR-186-5p is involved in responses to HIV.

We would like to emphasize several aspects of the study that open the door to future research:

1. We used stepped ultracentrifugation without density gradients because of the small sample volumes available. Although stepped ultracentrifugation remains a widely used method for EV enrichment ^43,90^, subsequent gradients or alternative isolation methods could be attempted with larger volume samples to increase purity in future. Possibly, our study overestimates the abundance of miRNAs in CVL EVs, and differential packaging into EVs and exRNPs is masked by contamination of our EV preps with exRNPs.
2. Our qPCR array approach and focus on miRNAs leaves room for additional work. While we are confident that our array captured most of the abundant miRNAs in CVL, sequencing short and longer RNAs could reveal additional markers.
3. The small number of subjects and the absence of obvious menstrual cycle in infected subjects precludes strong conclusions about EV or miRNA associations with either infection or the menstrual cycle. For example, we did not observe the expected increase in miR-451a or other red blood cell-specific miRNAs during menstruation. However, since only two animals showed evidence of cycling, experiments with more subjects and larger sample volumes are needed.
4. Our previous criticisms of miRNA functional studies ^91^ also apply to our results here. Additional work is needed to assess the potential of miR-186-5p to regulate retrovirus production at endogenous levels, for example by showing that it is present in active RNPs ^92^ and that it interacts directly with specific host or viral targets. However, it is also important to note that miR-186-5p could have therapeutic benefit even if it must be delivered at supraphysiologic concentrations. Finally, it is possible, but must be demonstrated, that miR-186-5p acts in a paracrine fashion via EV or exRNP shuttles.
5. We have investigated the effects of miR-186-5p only in monocyte-derived macrophages. We chose to begin with this cell type because of the abundance of miR-223 and the known role of macrophages in the epithelium. Other cell types should also be investigated.

Overall, the results presented here support further development of CVL and its constituents as a window into the health of the cervicovaginal compartment in retroviral infection and beyond. Furthermore, delivery of miR-186-5p could act to suppress retrovirus release.

## Supplementary Materials

Figure S1: Specimen Collection and Sample Processing Workflow, Figure S2: miRNA profile of CVL fractions, Table S1: Recovered volumes: CVL, Table S2: NTA dilution factors, CVL, Table S3: Pooling Strategy

## Author Contributions

Conceptualization, Zezhou Zhao, Dillon C. Muth, Grace V. Hancock, Kelly A. Metcalf Pate and Kenneth W. Witwer; Data curation, Zezhou Zhao and Kenneth W. Witwer; Formal analysis, Zezhou Zhao, Dillon C. Muth and Kenneth W. Witwer; Funding acquisition, Kenneth W. Witwer; Investigation, Zezhou Zhao, Dillon C. Muth, Kathleen Mulka, Zhaohao Liao and Bonita H. Powell; Methodology, Zezhou Zhao, Kathleen Mulka, Grace V. Hancock, Kelly A. Metcalf Pate and Kenneth W. Witwer; Project administration, Kenneth W. Witwer; Resources, Kelly A. Metcalf Pate and Kenneth W. Witwer; Supervision, Zhaohao Liao and Kenneth W. Witwer; Visualization, Zezhou Zhao, Dillon C. Muth and Kenneth W. Witwer; Writing – original draft, Kenneth W. Witwer; Writing – review & editing, Zezhou Zhao, Dillon C. Muth and Kenneth W. Witwer.

## Acknowledgments

The authors thank Robert Adams, Lauren Ostrenga, and Sarah Beck for contributions to these studies. The authors gratefully acknowledge the Oregon National Primate Research Center and David Erikson for hormone analyses and endocrinology advice and thank Barbara Smith of the JHU IBBS Microscope Facility for expert assistance with electron microscopy. Amanda Steele provided paid assistance with editing and organizing an early version of the manuscript.

## SUPPLEMENTAL MATERIALS

**Supplemental Figure S1.**
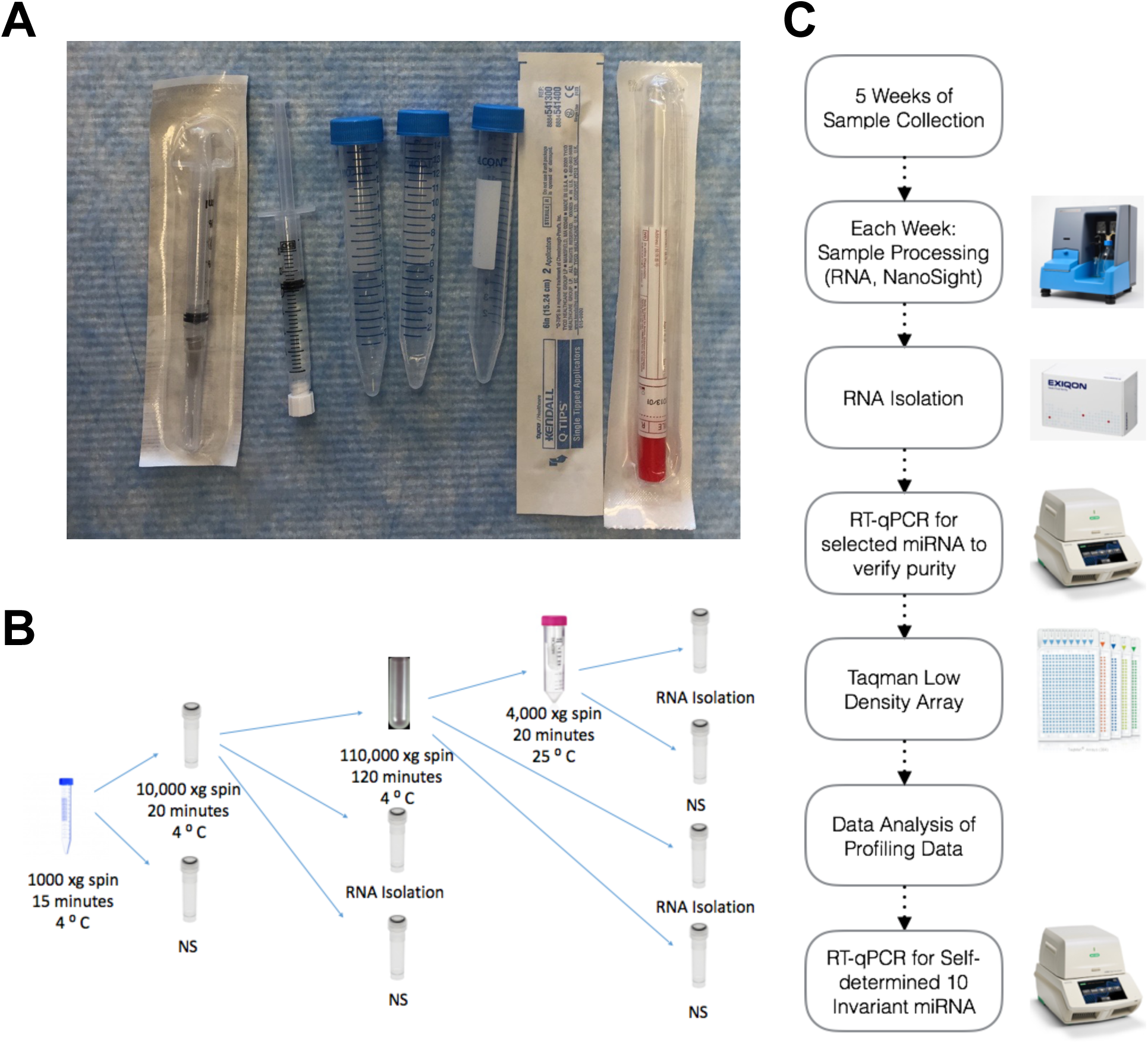
Collection Materials and Workflow

**Supplemental Figure S2.**
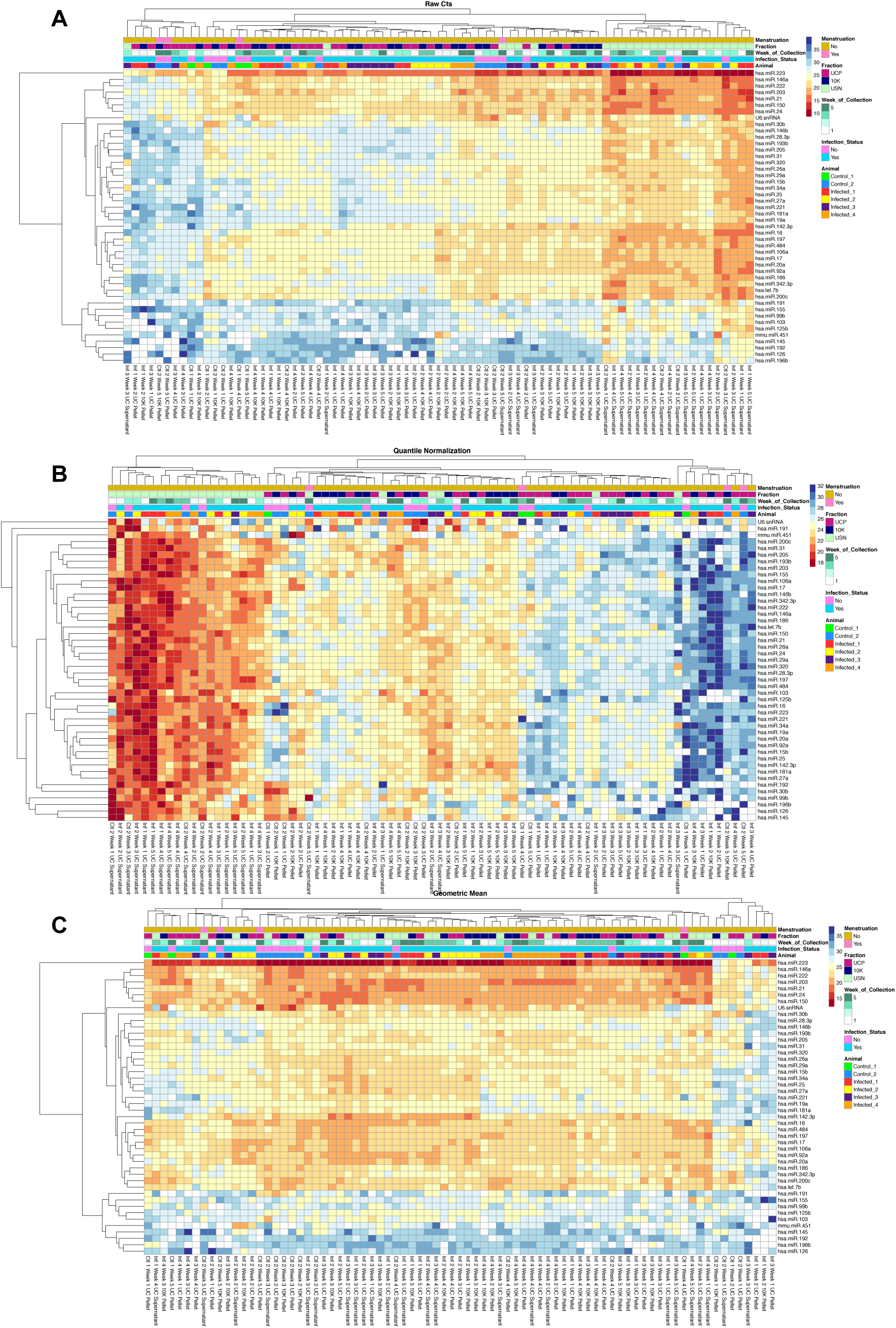

## SUPPLEMENTAL TABLES

**Supplemental Table 1.** Recovered volumes: CVL

| Subject | Week 1 | Week 2 | Week 3 | Week 4 | Week 5 |
| --- | --- | --- | --- | --- | --- |
| Control 1 | 1 mL | 2 mL | 1 mL | 1 mL | 1 mL |
| Control 2 | 3 mL | 0.7 mL | 2 mL | 0.5 mL | 1.25 mL |
| Infected 1 | 1 mL | 0.2 mL | 0.5 mL | 0.3 mL | 1.5 mL |
| Infected 2 | 1.5 mL | 1.5 mL | 2 mL | 0.8 mL | 1.75 mL |
| Infected 3 | 0.5 mL | 0.6 mL | 2.8 mL | 0.8 mL | 2 mL |
| Infected 4 | 0.8 mL | 0.3 mL | 0.6 mL | 0.3 mL | 1.5 mL |

**Supplemental Table 2.** NTA dilution factors, CVL

|  | Time point | Subject |  |  |  |  |  |
| --- | --- | --- | --- | --- | --- | --- | --- |
|  |  | Control 1 | Control 2 | Infected 1 | Infected 2 | Infected 3 | Infected 4 |
| CVL (UC Supernatant) | Week 1 | 1:25 | 1:50 | 1:25 | 1:25 | 1:25 | 1:25 |
|  | Week 2 | 1:100 | 1:5 | 1:25 | 1:10 | 1:5 | 1:5 |
|  | Week 3 | 1:10 | 1:5 | 1:5 | 1:5 | 1:10 | 1:10 |
|  | Week 4 | 1:10 | 1:5 | 1:10 | 1:5 | 1:10 | 1:10 |
|  | Week 5 | 1:5 | 1:10 | 1:10 | 1:10 | 1:5 | 1:5 |
| CVL (UC Pellet) | Week 1 | Neat | 1:5 | 1:5 | 1:5 | 1:5 | 1:5 |
|  | Week 2 | 1:10 | 1:10 | 1:5 | 1:5 | 1:5 | 1:5 |
|  | Week 3 | 1:5 | 1:5 | 1:5 | 1:5 | 1:5 | 1:5 |
|  | Week 4 | 1:5 | 1:5 | 1:5 | 1:5 | 1:5 | 1:5 |
|  | Week 5 | 1:5 | 1:5 | 1:5 | 1:5 | 1:5 | 1:5 |

Abbreviations: CVL = cervicovaginal lavage; UC = ultracentrifugation

**Supplemental Table 3.** Pooling strategy: Western blot

|  |  |
| --- | --- |
| **Pool 1 (healthy and infected)** | Healthy 2 Week 5, Infected 1 Week 2, Infected 3 Week 2, Infected 3 Week 5, Infected 2 Week 3, Infected 2 Week 4 |
| **Pool 2 (all healthy)** | Healthy 1 Week 4, Healthy 1 Week 5, Healthy 2 Week 2, Healthy 2 Week 3, Healthy 2 Week 4, Healthy 1 Week 2 |
| **Pool 3 (all infected)** | Infected 5 Week 2, Infected 3 Week 3, Infected 2 Week 4, Infected 4 Week 5, Infected 4 Week 3, Infected 1 Week 5 |

## REFERENCES

1. Hanson EK, Ballantyne J. Highly specific mRNA biomarkers for the identification of vaginal secretions in sexual assault investigations. Sci Justice. 2013;53(1):14–22. doi:10.1016/j.scijus.2012.03.007

2. Jakubowska J, MacIejewska A, Pawlowski R, Bielawski KP. MRNA profiling for vaginal fluid and menstrual blood identification. Forensic Sci Int Genet. 2013;7(2):272–278. doi:10.1016/j.fsigen.2012.11.005

3. Park JL, Kwon OH, Kim JH, et al. Identification of body fluid-specific DNA methylation markers for use in forensic science. Forensic Sci Int Genet. 2014;13:147–153. doi:10.1016/j.fsigen.2014.07.011

4. Hanson EK, Lubenow H, Ballantyne J. Identification of forensically relevant body fluids using a panel of differentially expressed microRNAs. Anal Biochem. 2009;387(2):303–314. doi:10.1016/j.ab.2009.01.037S0003-2697(09)00065-7 [pii]

5. Liu J, Sun H, Wang X, et al. Increased Exosomal MicroRNA-21 and MicroRNA-146a Levels in the Cervicovaginal Lavage Specimens of Patients with Cervical Cancer. Int J Mol Sci Int J Mol Sci. 2014;15:758–773. doi:10.3390/ijms15010758

6. Van Ostade X, Dom M, Tjalma W, Van Raemdonck G. Candidate biomarkers in the cervical vaginal fluid for the (self-)diagnosis of cervical precancer. Arch Gynecol Obstet. 2018;297(2):295–311. doi:10.1007/s00404-017-4587-2

7. Van Raemdonck GAA, Tjalma WAA, Coen EP, Depuydt CE, Van Ostade XWM. Identification of Protein Biomarkers for Cervical Cancer Using Human Cervicovaginal Fluid. PLoS One. 2014;9(9):e106488. doi:10.1371/journal.pone.0106488

8. Gravett MG, Thomas A, Schneider KA, et al. Proteomic Analysis of Cervical-Vaginal Fluid: Identification of Novel Biomarkers for Detection of Intra-amniotic Infection. J Proteome Res. 2007;6(1):89–96. doi:10.1021/pr060149v

9. Datcu R, Gesink D, Mulvad G, et al. Bacterial vaginosis diagnosed by analysis of first-void-urine specimens. J Clin Microbiol. 2014;52(1):218–225. doi:10.1128/JCM.02347-13

10. Srinivasan S, Morgan MT, Fiedler TL, et al. Metabolic signatures of bacterial vaginosis Supplementary file: Figure S1. MBio. 2015;6(2):1–16. doi:10.1128/mBio.00204-15

11. Zevin AS, Xie IY, Birse K, et al. Microbiome Composition and Function Drives Wound-Healing Impairment in the Female Genital Tract. PLoS Pathog. 2016;12(9):e1005889. doi:10.1371/journal.ppat.1005889

12. Boggiano C, Littman DR. HIV’s Vagina Travelogue. Immunity. 2007;26(2):145–147. doi:10.1016/j.immuni.2007.02.001

13. Patel M V., Ghosh M, Fahey J V., Ochsenbauer C, Rossoll RM, Wira CR. Innate Immunity in the Vagina (Part II): Anti-HIV Activity and Antiviral Content of Human Vaginal Secretions. Am J Reprod Immunol. 2014;72(1):22–33. doi:10.1111/aji.12218

14. Burgener A, Boutilier J, Wachihi C, et al. Identification of differentially expressed proteins in the cervical mucosa of HIV-1-resistant sex workers. J Proteome Res. 2008;7(10):4446–4454. doi:10.1021/pr800406r

15. Benki S, Mostad SB, Richardson BA, Mandaliya K, Kreiss JK, Overbaugh J. Increased levels of HIV-1-infected cells in endocervical secretions after the luteinizing hormone surge. J Acquir Immune Defic Syndr. 2008;47(5):529–534. doi:10.1097/QAI.0b013e318165b952

16. Zara F, Nappi RE, Brerra R, Migliavacca R, Maserati R, Spinillo A. Markers of local immunity in cervico-vaginal secretions of HIV infected women: implications for HIV shedding. Sex Transm Infect. 2004;80(2):108–112. doi:10.1136/sti.2003.005157

17. Gardella B, Roccio M, Maccabruni A, et al. HIV shedding in cervico-vaginal secretions in pregnant women. Curr HIV Res. 2011;9(5):313–320. doi:Abs: CHIVR-162 [pii]

18. Seaton KE, Ballweber L, Lan A, et al. HIV-1 specific IgA detected in vaginal secretions of HIV uninfected women participating in a microbicide trial in Southern Africa are primarily directed toward gp120 and gp140 specificities. PLoS One. 2014;9(7). doi:10.1371/journal.pone.0101863

19. Ghosh M, Fahey J V., Shen Z, et al. Anti-HIV activity in cervical-vaginal secretions from HIV-Positive and -Negative women correlate with innate antimicrobial levels and IgG antibodies. PLoS One. 2010;5(6). doi:10.1371/journal.pone.0011366

20. Clemetson DB, Moss GB, Willerford DM, et al. Detection of HIV DNA in cervical and vaginal secretions. Prevalence and correlates among women in Nairobi, Kenya. Jama. 1993;269(22):2860–2864. doi:10.1016/0020-7292(94)90090-6

21. Burgener A, Rahman S, Ahmad R, et al. Comprehensive proteomic study identifies serpin and cystatin antiproteases as novel correlates of HIV-1 resistance in the cervicovaginal mucosa of female sex workers. J Proteome Res. 2011;10(11):5139–5149. doi:10.1021/pr200596r

22. Drannik AG, Nag K, Yao XD, et al. Anti-HIV-1 Activity of Elafin Depends on Its Nuclear Localization and Altered Innate Immune Activation in Female Genital Epithelial Cells. PLoS One. 2012;7(12). doi:10.1371/journal.pone.0052738

23. Yáñez-Mó M, Siljander PR-M, Andreu Z, et al. Biological properties of extracellular vesicles and their physiological functions. J Extracell vesicles. 2015;4:27066. doi:10.3402/jev.v4.27066

24. Théry C, Witwer KW, Aikawa E, et al. Minimal information for studies of extracellular vesicles 2018 (MISEV2018): a position statement of the International Society for Extracellular Vesicles and update of the MISEV2014 guidelines. J Extracell Vesicles. 2018;7(1). doi:10.1080/20013078.2018.1535750

25. Witwer KW, Théry C. Extracellular vesicles or exosomes? On primacy, precision, and popularity influencing a choice of nomenclature. J Extracell Vesicles. 2019;8(1):1648167. doi:10.1080/20013078.2019.1648167

26. Meehan B, Rak J, Di Vizio D. Oncosomes - large and small: what are they, where they came from? J Extracell vesicles. 2016;5:33109.

27. György B, Hung ME, Breakefield XO, Leonard JN. Therapeutic applications of extracellular vesicles: clinical promise and open questions. Annu Rev Pharmacol Toxicol. 2015;55:439–464. doi:10.1146/annurev-pharmtox-010814-124630

28. Witwer KW, Buzás EI, Bemis LT, et al. Standardization of sample collection, isolation and analysis methods in extracellular vesicle research. J Extracell vesicles. 2013;2:1–25. doi:10.3402/jev.v2i0.20360

29. Smith JA, Daniel R. Human vaginal fluid contains exosomes that have an inhibitory effect on an early step of the HIV-1 life cycle. AIDS. August 2016. doi:10.1097/QAD.0000000000001236

30. Muth DC, McAlexander MA, Ostrenga LJ, et al. Potential role of cervicovaginal extracellular particles in diagnosis of endometriosis. Bmc Vet Res. 2015. doi:10.1186/s12917-015-0513-7

31. Sergeeva AM, Pinzon Restrepo N, Seitz H. Quantitative aspects of RNA silencing in metazoans. Biochem. 2013;78(6):613–626. doi:10.1134/S0006297913060072BCM78060795 [pii]

32. Bartel DP. MicroRNAs: target recognition and regulatory functions. Cell. 2009;136(2):215–233. doi:S0092-8674(09)00008-7 [pii]10.1016/j.cell.2009.01.002

33. Valadi H, Ekström K, Bossios A, Sjöstrand M, Lee JJ, Lötvall JO. Exosome-mediated transfer of mRNAs and microRNAs is a novel mechanism of genetic exchange between cells. Nat Cell Biol. 2007;9(6):654–659. doi:10.1038/ncb1596

34. Aliotta JM, Sanchez-Guijo FM, Dooner GJ, et al. Alteration of marrow cell gene expression, protein production, and engraftment into lung by lung-derived microvesicles: a novel mechanism for phenotype modulation. Stem Cells. 2007;25(9):2245–2256. doi:10.1634/stemcells.2007-0128

35. Baj-Krzyworzeka M, Szatanek R, Węglarczyk K, et al. Tumour-derived microvesicles carry several surface determinants and mRNA of tumour cells and transfer some of these determinants to monocytes. Cancer Immunol Immunother. 2006;55(7):808–818. doi:10.1007/s00262-005-0075-9

36. Mateescu B, Kowal EJK, van Balkom BWM, et al. Obstacles and opportunities in the functional analysis of extracellular vesicle RNA - an ISEV position paper. J Extracell vesicles. 2017;6(1):1286095. doi:10.1080/20013078.2017.1286095

37. Turchinovich A, Weiz L, Langheinz A, Burwinkel B. Characterization of extracellular circulating microRNA. Nucleic Acids Res. 2011;39(16):7223–7233. doi:gkr254 [pii]10.1093/nar/gkr254

38. Arroyo JD, Chevillet JR, Kroh EM, et al. Argonaute2 complexes carry a population of circulating microRNAs independent of vesicles in human plasma. Proc Natl Acad Sci U S A. 2011;108(12):5003–5008. doi:1019055108 [pii]10.1073/pnas.1019055108

39. Witwer KW, Sarbanes SL, Liu J, Clements JE. A plasma microRNA signature of acute lentiviral infection: biomarkers of CNS disease. AIDS. 2011;204(7):1104–1114. doi:10.1097/QAD.0b013e32834b95bf

40. Zubakov D, Boersma AW, Choi Y, van Kuijk PF, Wiemer EA, Kayser M. MicroRNA markers for forensic body fluid identification obtained from microarray screening and quantitative RT-PCR confirmation. Int J Leg Med. 2010;124(3):217–226. doi:10.1007/s00414-009-0402-3

41. Seashols-Williams S, Lewis C, Calloway C, et al. High-throughput miRNA sequencing and identification of biomarkers for forensically relevant biological fluids. Electrophoresis. August 2016. doi:10.1002/elps.201600258

42. Rahman S, Rabbani R, Wachihi C, et al. Mucosal serpin A1 and A3 levels in HIV highly exposed sero-negative women are affected by the menstrual cycle and hormonal contraceptives but are independent of epidemiological confounders. Am J Reprod Immunol. 2013;69(1):64–72. doi:10.1111/aji.12014

43. Thery C, Amigorena S, Raposo G, Clayton A. Isolation and characterization of exosomes from cell culture supernatants and biological fluids. Curr Protoc Cell Biol. 2006;Chapter 3:Unit 3 22. doi:10.1002/0471143030.cb0322s30

44. McAlexander MA, Phillips MJ, Witwer KW. Comparison of methods for miRNA extraction from plasma and quantitative recovery of RNA from cerebrospinal fluid. Front Genet. 2013;4(MAY):83. doi:10.3389/fgene.2013.00083

45. Chen C, Ridzon DA, Broomer AJ, et al. Real-time quantification of microRNAs by stem-loop RT-PCR. Nucleic Acids Res. 2005;33(20):e179. doi:33/20/e179 [pii]10.1093/nar/gni178

46. Witwer KW, Sarbanes SL, Liu J, Clements JE. A plasma microRNA signature of acute lentiviral infection: biomarkers of central nervous system disease. AIDS. 2011;25(17):2057–2067. doi:10.1097/QAD.0b013e32834b95bf

47. Clough E, Barrett T. The Gene Expression Omnibus Database. Methods Mol Biol. 2016;1418:93–110. doi:10.1007/978-1-4939-3578-9_5

48. Van Deun J, Mestdagh P, Agostinis P, et al. EV-TRACK: transparent reporting and centralizing knowledge in extracellular vesicle research. Nat Methods. 2017;14(3):228–232. doi:10.1038/nmeth.4185

49. Biton M, Levin A, Slyper M, et al. Epithelial microRNAs regulate gut mucosal immunity via epithelium–T cell crosstalk. Nat Immunol. 2011;12(3):239–246. doi:10.1038/ni.1994

50. Jurcevic S, Olsson B, Klinga-Levan K. MicroRNA expression in human endometrial adenocarcinoma. Cancer Cell Int. 2014;14(1):1–8. doi:10.1186/s12935-014-0088-6

51. Jayaraman M, Radhakrishnan R, Mathews CA, et al. Identification of novel diagnostic and prognostic miRNA signatures in endometrial cancer. Genes Cancer. 2017;8(5-6):566–576. doi:10.18632/genesandcancer.144

52. Cosar E, Mamillapalli R, Ersoy GS, Cho SY, Seifer B, Taylor HS. Serum microRNAs as diagnostic markers of endometriosis: a comprehensive array-based analysis. Fertil Steril. 2016;106(2):402–409. doi:10.1016/j.fertnstert.2016.04.013

53. Schwarzenbach H, da Silva AM, Calin G, Pantel K. Data Normalization Strategies for MicroRNA Quantification. Clin Chem. 2015;61(11):1333–1342. doi:10.1373/clinchem.2015.239459

54. Nguyen HPT, Simpson RJ, Salamonsen LA, Greening DW. Extracellular Vesicles in the Intrauterine Environment: Challenges and Potential Functions. Biol Reprod. 2016;95(5):109–109. doi:10.1095/biolreprod.116.143503

55. Campoy I, Lanau L, Altadill T, et al. Exosome-like vesicles in uterine aspirates: a comparison of ultracentrifugation-based isolation protocols. J Transl Med. 2016;14(1):180. doi:10.1186/s12967-016-0935-4

56. Zegels G, Aa G, Raemdonck V, et al. Comprehensive proteomic analysis of human cervical-vaginal fluid using colposcopy samples. Proteome Sci. 2009;7(7). doi:10.1186/1477-5956-7-17

57. Gravett MG, Thomas A, Schneider KA, et al. PROTEOMIC ANALYSIS OF CERVICAL-VAGINAL FLUID: IDENTIFICATION OF NOVEL BIOMARKERS FOR DETECTION OF INTRA-AMNIOTIC INFECTION. doi:10.1021/pr060149v

58. Arroyo JD, Chevillet JR, Kroh EM, et al. Argonaute2 complexes carry a population of circulating microRNAs independent of vesicles in human plasma. Proc Natl Acad Sci U S A. 2011;108(12):5003–5008. doi:10.1073/pnas.1019055108

59. Guo Z, Maki M, Ding R, Yang Y, Zhang B, Xiong L. Genome-wide survey of tissue-specific microRNA and transcription factor regulatory networks in 12 tissues. Sci Rep. 2014;4:1–9. doi:10.1038/srep05150

60. Panwar B, Omenn GS, Guan Y. miRmine: A Database of Human miRNA Expression Profiles. Bioinformatics. 2017;33(10):btx019. doi:10.1093/bioinformatics/btx019

61. Pritchard CC, Kroh E, Wood B, et al. Blood cell origin of circulating microRNAs: a cautionary note for cancer biomarker studies. Cancer Prev Res. 2012;5(3):492–497. doi:1940-6207.CAPR-11-0370 [pii]10.1158/1940-6207.CAPR-11-0370

62. Luo P, Wang Q, Ye Y, et al. MiR-223-3p functions as a tumor suppressor in lung squamous cell carcinoma by miR-223-3p-mutant p53 regulatory feedback loop. J Exp Clin Cancer Res. 2019;38(1). doi:10.1186/s13046-019-1079-1

63. Ji Q, Xu X, Song Q, et al. miR-223-3p Inhibits Human Osteosarcoma Metastasis and Progression by Directly Targeting CDH6. Mol Ther. 2018;26:1299–1312. doi:10.1016/j.ymthe.2018.03.009

64. Lawrence M JY. miR-203 Functions as a Tumor Suppressor by Inhibiting Epithelial to Mesenchymal Transition in Ovarian Cancer. J Cancer Sci Ther. 2015;07(02). doi:10.4172/1948-5956.1000322

65. Deng B, Wang B, Fang J, et al. MiRNA-203 suppresses cell proliferation, migration and invasion in colorectal cancer via targeting of EIF5A2. Sci Rep. 2016;6. doi:10.1038/srep28301

66. Wang S, Zhang R, Claret FX, Yang H. Involvement of microRNA-24 and DNA methylation in resistance of nasopharyngeal carcinoma to ionizing radiation. Mol Cancer Ther. 2014;13(12):3163–3174. doi:10.1158/1535-7163.MCT-14-0317

67. Fang ZH, Wang SL, Zhao JT, et al. MIR-150 exerts antileukemia activity in vitro and in vivo through regulating genes in multiple pathways. Cell Death Dis. 2016;7(9). doi:10.1038/cddis.2016.256

68. Ito M, Teshima K, Ikeda S, et al. MicroRNA-150 inhibits tumor invasion and metastasis by targeting the chemokine receptor CCR6, in advanced cutaneous T-cell lymphoma. Blood. 2014;123(10):1499–1511. doi:10.1182/blood-2013-09-527739

69. Huang J, Wang F, Argyris E, et al. Cellular microRNAs contribute to HIV-1 latency in resting primary CD4+ T lymphocytes. Nat Med. 2007;13(10):1241–1247. doi:nm1639 [pii]10.1038/nm1639

70. Swaminathan S, Murray DD, Kelleher AD. miRNAs and HIV: unforeseen determinants of host-pathogen interaction. Immunol Rev. 2013;254(1):265–280. doi:10.1111/imr.12077

71. Sisk JM, Witwer KW, Tarwater PM, Clements JE. SIV replication is directly downregulated by four antiviral miRNAs. Retrovirology. 2013;10(1):95. doi:1742-4690-10-95 [pii]10.1186/1742-4690-10-95

72. Wang X, Ye L, Zhou Y, Liu MQ, Zhou DJ, Ho WZ. Inhibition of anti-HIV microRNA expression: a mechanism for opioid-mediated enhancement of HIV infection of monocytes. Am J Pathol. 2011;178(1):41–47. doi:S0002-9440(10)00089-1 [pii]10.1016/j.ajpath.2010.11.042

73. Swaminathan S, Suzuki K, Seddiki N, et al. Differential regulation of the Let-7 family of microRNAs in CD4+ T cells alters IL-10 expression. J Immunol. 2012;188(12):6238–6246. doi:jimmunol.1101196 [pii]10.4049/jimmunol.1101196

74. Klase Z, Kale P, Winograd R, et al. HIV-1 TAR element is processed by Dicer to yield a viral micro-RNA involved in chromatin remodeling of the viral LTR. BMC Mol Biol. 2007;8:63. doi:1471-2199-8-63 [pii]10.1186/1471-2199-8-63

75. Wagschal A, Rousset E, Basavarajaiah P, et al. Microprocessor, Setx, Xrn2, and Rrp6 co-operate to induce premature termination of transcription by RNAPII. Cell. 2012;150(6):1147–1157. doi:10.1016/j.cell.2012.08.004S0092-8674(12)00999-3 [pii]

76. Whisnant AW, Bogerd HP, Flores O, et al. In-Depth Analysis of the Interaction of HIV-1 with Cellular microRNA Biogenesis and Effector Mechanisms. MBio. 2013;4(2). doi:10.1128/mBio.00193-13e00193-13 [pii]mBio.00193-13 [pii]

77. Sung TL, Rice AP. miR-198 inhibits HIV-1 gene expression and replication in monocytes and its mechanism of action appears to involve repression of cyclin T1. PLoS Pathog. 2009;5(1):e1000263. doi:10.1371/journal.ppat.1000263

78. Hsu SD, Tseng YT, Shrestha S, et al. miRTarBase update 2014: an information resource for experimentally validated miRNA-target interactions. Nucleic Acids Res. 2014;42(Database issue):D78–85. doi:10.1093/nar/gkt1266

79. Hsu SD, Lin FM, Wu WY, et al. miRTarBase: a database curates experimentally validated microRNA-target interactions. Nucleic Acids Res. 2011;39(Database issue):D163–9. doi:10.1093/nar/gkq1107

80. Vergoulis T, Vlachos IS, Alexiou P, et al. TarBase 6.0: capturing the exponential growth of miRNA targets with experimental support. Nucleic Acids Res. 2011. doi:gkr1161 [pii]10.1093/nar/gkr1161

81. Vlachos IS, Paraskevopoulou MD, Karagkouni D, et al. DIANA-TarBase v7.0: indexing more than half a million experimentally supported miRNA:mRNA interactions. Nucleic Acids Res. 2015;43(Database issue):D153–9. doi:10.1093/nar/gku1215

82. Lagana A, Forte S, Giudice A, et al. miRo: a miRNA knowledge base. Database. 2009;2009(0):bap008–bap008. doi:10.1093/database/bap008

83. Papadopoulos GL, Alexiou P, Maragkakis M, Reczko M, Hatzigeorgiou AG. DIANA-mirPath: Integrating human and mouse microRNAs in pathways. Bioinformatics. 2009;25(15):1991–1993. doi:10.1093/bioinformatics/btp299

84. Vlachos IS, Zagganas K, Paraskevopoulou MD, et al. DIANA-miRPath v3.0: deciphering microRNA function with experimental support. Nucleic Acids Res. 2015;43(W1):W460–W466. doi:10.1093/nar/gkv403

85. Myatt SS, Wang J, Monteiro LJ, et al. Definition of microRNAs that repress expression of the tumor suppressor gene FOXO1 in endometrial cancer. Cancer Res. 2010;70(1):367–377. doi:10.1158/0008-5472.CAN-09-1891

86. Zhou L, Qi X, Potashkin JA, Abdul-Karim FW, Gorodeski GI. MicroRNAs miR-186 and miR-150 down-regulate expression of the pro-apoptotic purinergic P2X7 receptor by activation of instability sites at the 3’-untranslated region of the gene that decrease steady-state levels of the transcript. J Biol Chem. 2008;283(42):28274–28286. doi:10.1074/jbc.M802663200

87. Niinuma T, Kai M, Kitajima H, et al. Downregulation of miR-186 is associated with metastatic recurrence of gastrointestinal stromal tumors. Oncol Lett. 2017;14(5):5703–5710. doi:10.3892/ol.2017.6911

88. Paraskevopoulou MD, Vlachos IS, Hatzigeorgiou AG. DIANA-TarBase and DIANA Suite Tools: Studying Experimentally Supported microRNA Targets. Curr Protoc Bioinforma. 2016;55(1):12.14.1-12.14.18. doi:10.1002/cpbi.12

89. Ji Diana Lee Y, Kim V, Muth DC, Witwer KW. Validated MicroRNA Target Databases: An Evaluation. Drug Dev Res. 2015;76(7):389–396. doi:10.1002/ddr.21278

90. Gardiner C, Vizio D Di, Sahoo S, et al. Techniques used for the isolation and characterization of extracellular vesicles: results of a worldwide survey. J Extracell Vesicles. 2016;5(0). doi:10.3402/jev.v5.32945

91. Witwer KW, Halushka MK. Towards the Promise of microRNAs - Enhancing reproducibility and rigor in microRNA research. RNA Biol. September 2016:0. doi:10.1080/15476286.2016.1236172

92. La Rocca G, Olejniczak SH, Gonzalez AJ, et al. In vivo, Argonaute-bound microRNAs exist predominantly in a reservoir of low molecular weight complexes not associated with mRNA. Proc Natl Acad Sci U S A. 2015;112(3):767–772. doi:10.1073/pnas.1424217112

